# A synthetic biology platform for the reconstitution and mechanistic dissection of LINC complex assembly

**DOI:** 10.1101/415745

**Authors:** Sagardip Majumder, Patrick T. Willey, Maxwell S. DeNies, Allen P. Liu, G.W. Gant Luxton

## Abstract

The linker of nucleoskeleton and cytoskeleton (LINC) is a conserved nuclear envelope-spanning molecular bridge that is responsible for the mechanical integration of the nucleus with the cytoskeleton. LINC complexes are formed by a transluminal interaction between the outer and inner nuclear membrane KASH and SUN proteins, respectively. Despite recent structural insights, our mechanistic understanding of LINC complex assembly remains limited by the lack of an experimental system for its *in vitro* reconstitution and manipulation. Here, we describe artificial nuclear membranes (ANMs) as a synthetic biology platform based on mammalian cell-free expression for the rapid reconstitution of SUN proteins in supported lipid bilayers. We demonstrate that SUN1 and SUN2 are oriented in ANMs with solvent-exposed C-terminal KASH-binding SUN domains. We also find that SUN2 possesses a single transmembrane domain, while SUN1 possesses three. Finally, SUN protein-containing ANMs bind synthetic KASH peptides, thereby reconstituting the LINC complex core. This work represents the first *in vitro* reconstitution of KASH-binding SUN proteins in supported lipid bilayers using cell-free expression, which will be invaluable for testing proposed models of LINC complex assembly and its regulation.

## INTRODUCTION

Eukaryotic cells are defined by the presence of a genome-containing nucleus, the boundary of which is delineated by the nuclear envelope (NE), a specialized subdomain of the endoplasmic reticulum (ER) (Kite, 1913). The NE consists of concentric inner and outer nuclear membranes (INM and ONM, respectively) separated by a ~30-50 nm perinuclear space (PNS) that is contiguous with the ER lumen (Watson, 1959). While the ONM is an extension of the ER, a unique subset of proteins resides in the INM that interact with the nuclear lamina and chromatin within the nucleoplasm (Burke and Stewart, 2014).

Fusion between the INM and ONM creates numerous aqueous channels throughout the NE that are occupied by nuclear pore complexes (NPCs), which are the primary sites of molecular exchange between the cytoplasm and nucleoplasm (Knockenhauer and Schwartz, 2016, Otsuka and Ellenberg, 2018). However, mechanical forces generated by the cytoskeleton within the cytoplasm can also be sensed and transmitted across the NE and into the nucleoplasm by LINC complexes (Alam et al., 2016, Brosig et al., 2010, Guilluy et al., 2014, Lombardi et al., 2011, Tajik et al., 2016). These evolutionarily conserved NE-spanning molecular bridges mediate several fundamental cellular processes including DNA damage repair, meiotic chromosome pairing, mechano-regulation of gene expression, and nuclear positioning (Alam et al., 2016, Chang et al., 2015, Meinke and Schirmer, 2015, Tapley and Starr, 2013). Consistent with their central role in cellular function is a growing list of genetic mutations in LINC complex proteins associated with human diseases such as aging-related hearing loss, ataxia, and muscular dystrophy (Horn, 2014, Janin et al., 2017).

LINC complexes are formed by the transluminal interactions of the ONM Klarsicht/ANC-1/SYNE homology (KASH) proteins and the INM Sad1/UNC-84 (SUN) proteins (Crisp et al., 2006). The divergent cytoskeletal-binding cytoplasmic domain of KASH proteins largely consists of spectrin repeats or coiled-coils (CCs) (Luxton and Starr, 2014, Meinke and Schirmer, 2015), while their C-termini contain the conserved nuclear envelope targeting KASH domain composed of a transmembrane domain (TMD) followed by the ~10-32 residue luminal KASH peptide (Starr and Han, 2002). Within the PNS, the C-terminal SUN domain of SUN proteins interacts with KASH peptides (Malone et al., 1999), whereas their divergent N-termini reside within the nucleoplasm where they interact with A-type lamins, chromatin, as well as other INM proteins (Chang et al., 2015). Mammals encode six KASH proteins (nesprins 1-4, lymphocyte-restricted membrane protein, and KASH5) and five SUN proteins (SUN1-5) (Meinke and Schirmer, 2015, Stewart and Burke, 2014). The expression of SUN3-5 is testis-specific and their ability to assemble into functional LINC complexes remains unclear (Sosa et al., 2013). In contrast, SUN1 and SUN2 are both widely expressed in somatic cells and interact with all known KASH proteins (Stewart-Hutchinson et al., 2008).

Groundbreaking *in vitro* studies have provided critical insights into the mechanism of LINC complex assembly. In key papers describing its crystal structure, the Kutay, Schwartz, and Wang laboratories showed that SUN2 homo-trimerizes and that a short preceding CC was necessary and sufficient for homo-oligomerization (Sosa et al., 2012, Zhou et al., 2012). In addition, Kutay and Schwartz revealed that SUN2 homo-oligomerization was required for KASH peptide binding, which further stabilized the homo-trimer (Sosa et al., 2012). In an extension of this earlier work, the Feng laboratory suggested that the CC-containing region of the SUN2 luminal domain should be viewed as a pair of CCs, which potentially influence the monomer-trimer equilibrium of SUN2 (Nie et al., 2016). Using fluorescence fluctuation spectroscopy, an imaging-based technique that enables the quantification of protein oligomerization *in vivo* (Slaughter and Li, 2010), we recently confirmed that the luminal domain of SUN2 homo-trimerizes within the NE of living cells (Hennen et al., 2017, Hennen et al., 2018).

While SUN2 and SUN1 share a high level of sequence similarity (~65% identity between mouse SUN2 and SUN1), display similar affinity for the nesprin-2 KASH peptide (Ostlund et al., 2009), and perform several redundant cellular functions (i.e. DNA damage repair (Lei et al., 2012) and sub-synaptic nuclear anchorage in skeletal muscle (Lei et al., 2009)), we found that the homo-oligomerization of the SUN1 luminal domain within the NE was not limited to a homo-trimer (Hennen et al., 2018). Taken together, these results suggest that LINC complexes containing SUN1 assemble via a distinct mechanism from those containing SUN2, which may explain the specific requirement for SUN1 during meiotic chromosome pairing (Ding et al., 2007, Horn et al., 2013) as well as NPC assembly and distribution throughout the NE (Liu et al., 2007, Lu et al., 2008, Talamas and Hetzer, 2011).

To date, our understanding of the mechanisms underlying the differential assembly and function of SUN1-and SUN2-containing LINC complexes remains limited by the fact that the results described above were all generated using soluble fragments of the luminal domains of SUN1 and SUN2, not full-length SUN proteins within the context of a lipid bilayer. To begin to overcome this limitation, we describe here the development of artificial nuclear membranes (ANMs) as a simple bottom-up synthetic biology platform based on mammalian cell-free expression (CFE) for the rapid reconstitution and mechanistic dissection of LINC complex assembly composed of full-length SUN proteins.

## RESULTS

### Synthesis of SUN1 and SUN2 using mammalian CFE

CFE systems enable the synthesis of proteins of interest encoded by cDNA constructs in a one-pot transcription-translation reaction (Murray and Baliga, 2013, Rosenblum and Cooperman, 2014). Recent studies demonstrate the successful reconstitution of membrane proteins in ER-derived microsomes and supported lipid bilayers following CFE in eukaryotic cell extracts (Quast et al., 2015, Quast et al., 2016, Zemella et al., 2017). The use of CFE for mechanistic structure/function-based studies of membrane proteins offers several important advantages over more conventional cell-based methods where membrane proteins are over-expressed in and purified from heterologous systems (Kai et al., 2015, Schwarz et al., 2008). Specifically, CFE systems significantly reduce the risk of membrane protein denaturation, as newly synthesized membrane proteins are directly inserted into natural ER-based lipid bilayers in the absence of detergents, thus allowing for their proper folding. In addition, CFE is significantly more robust and efficient than the time-consuming process of cell-based expression followed by purification and reconstitution.

To investigate the feasibility of using CFE to reconstitute LINC complexes, we tested the ability of our previously described CFE system generated using lysates prepared from HeLa S3 cells grown in suspension (Ho et al., 2015, Ho et al., 2017) to synthesize full-length (FL) mouse SUN1 and SUN2 in the absence of exogenously supplied artificial membranes (Figure 1A). To do this, we sub-cloned previously characterized cDNA constructs encoding EGFP-tagged SUN1^FL^ and SUN2^FL^ (Luxton et al., 2010, Ostlund et al., 2009) behind the T7 promoter in the pT7CFE1-CHis vector for mammalian CFE to generate EGFP-SUN1^FL^-His_6_ and EGFP-SUN2^FL^-His_6_, respectively (Figure 1B). These constructs were then added separately to CFE reactions containing T7 RNA polymerase and incubated at 32°C in a 96 well plate. Protein production was monitored in a plate reader by quantifying bulk EGFP fluorescence for 2 hours, which revealed robust synthesis of both EGFP-SUN1^FL^-His_6_ and EGFP-SUN2^FL^-His_6_ above background levels (Figure 1C). Western blot analysis of these CFE reactions using anti-EGFP antibodies demonstrated single bands at molecular weights consistent with the synthesis of FL of EGFP-SUN1^FL^-His_6_ (MW ~131 kDa) and EGFP-SUN2^FL^-His_6_ (MW ~110 kDa) (Figure 1D). To determine if EGFP-SUN1^FL^-His_6_ and EGFP-SUN2^FL^-His_6_ were membrane-associated, we stained CFE reactions expressing either protein with the lipophilic membrane stain (1,1’-Dioctadecyl-3,3,3’,3’-Tetramethylindocarbocyanine Perchlorate (DiI). Confocal images taken away from the coverslip of these stained CFE reactions revealed punctae of EGFP and DiI fluorescence; however, we were unable to observe a clear co-localization between these punctae due to the speed of their diffusion and the temporal limitation of our microscope set-up (Fig. S1A). Nevertheless, co-localization between EGFP and DiI was observed in punctae that had settled down onto the coverslip suggesting that EGFP-SUN1^FL^-His_6_ and EGFP-SUN2^FL^-His_6_ are inserted into the ER-derived microsomes present within the HeLa S3 cell extracts used for their CFE (Fig. S1B). Taken together, these results show that our mammalian CFE system can be used for the efficient synthesis of FL SUN1 and SUN2.

**Figure 1:**
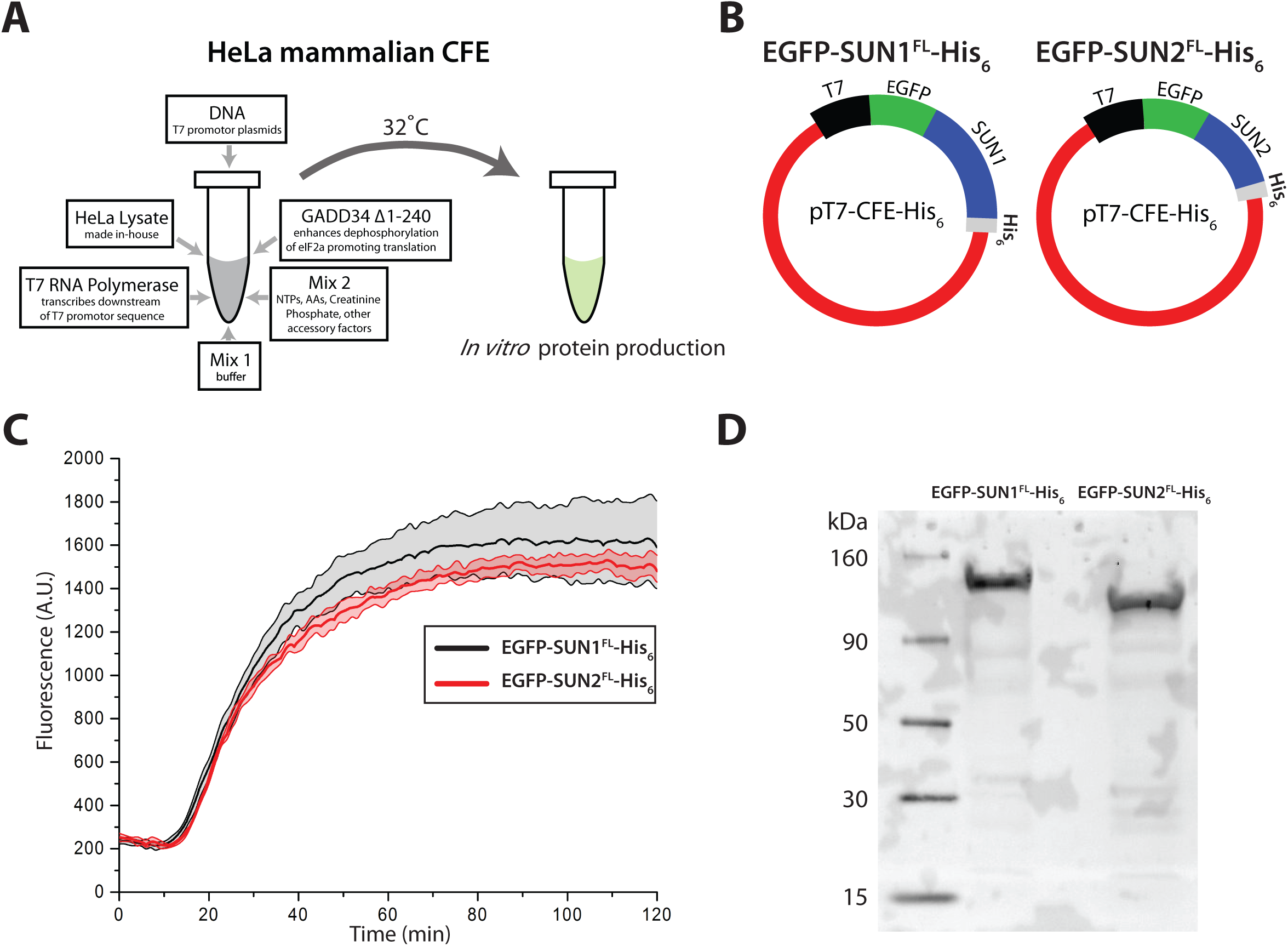
Synthesis of FL SUN1 and SUN2 using mammalian CFE. **A)** Schematic of the HeLa CFE system used in this work. **B)** Illustration of the pT7-CFE-His_6_ constructs encoding EGFP-SUN1^FL^-His_6_ and EGFP-SUN2^FL^-His_6_. **C)** Plot of the kinetics of EGFP-SUN1^FL^-His_6_ and EGFP-SUN2^FL^-His_6_ synthesis in HeLa CFE reactions as a read-out of EGFP fluorescence over time. Error bands indicate the standard deviation calculated from 3 independent experiments. **D)** Western blot of CFE-synthesized EGFP-SUN1^FL^-His_6_ and EGFP-SUN2^FL^-His_6_ probed with an anti-EGFP antibody.

### Insertion of CFE-generated SUN1 and SUN2 into ANMs

Our next step towards LINC complex reconstitution was to determine if our CFE synthesized SUN proteins could be inserted into exogenously provided artificial lipid bilayer membranes. Recently, we demonstrated the successful insertion of the bacterial mechanosensitive channel of large conductance, MscL, into artificial lipid vesicles following its synthesis in bacterial or mammalian CFE reactions (Ho et al., 2017, Majumder et al., 2017). Here, we followed a similar strategy; however, to facilitate the isolation of functional SUN protein inserted into artificial lipid bilayer membranes, we used supported lipid bilayer with excess membrane reservoir (SUPER) templates. Initially developed for *in vitro* studies of protein-mediated membrane fission (Pucadyil and Schmid, 2008, Pucadyil and Schmid, 2010), SUPER templates allow for the reconstitution of excess lipid bilayer membranes on 5 μm diameter silica beads due to the fusion of small unilamellar vesicles (SUVs) containing negatively charged lipids under conditions of high-ionic strength (Figure 2A). The lipid composition used for the SUVs in this study was designed to closely mimic that of the INM (van Meer et al., 2008). Following their assembly, the SUPER templates were added to a CFE reaction containing synthesized SUN proteins.

**Figure 2:**
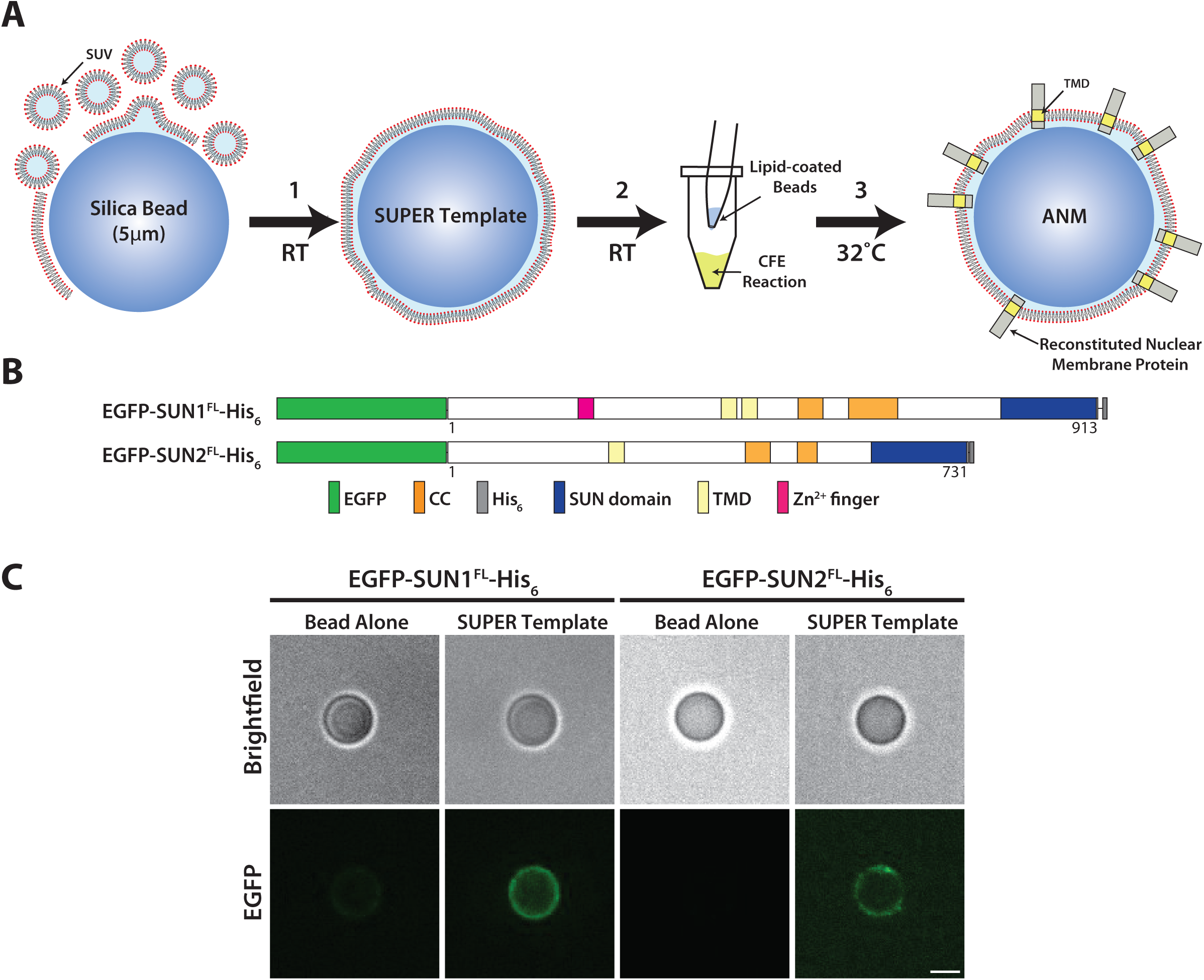
Reconstitution of CFE-synthesized FL SUN1 and SUN2 in ANMs. **A)** Schematic of the process of generating ANMs with inserted CFE-synthesized membrane proteins. **B)** Illustrations of the constructs used in this figure. **C)** Representative images of silica beads or SUPER templates incubated in CFE reactions of the indicated constructs. Scale bar: 5 μm.

We observed the successful association of both EGFP-SUN1^FL^-His_6_ and EGFP-SUN2^FL^-His_6_ with SUPER templates incubated in their CFE reactions (Figures 2B-C). Analysis of the protein domain architectures of SUN1^FL^ and SUN2^FL^ using the simple modular architecture research tool (SMART) (Schultz et al., 2000) predicted the existence of one and two TMDs, respectively. Importantly, neither construct associated with silica beads in the absence of SUPER template strongly suggesting the specific insertion of EGFP-SUN1^FL^-His_6_ and EGFP-SUN2^FL^-His_6_ into the lipid bilayer membrane of the SUPER template and the assembly of SUN protein-containing ANMs. Although we limit our use of ANMs in this work to reconstitute INM proteins, they could easily be used to reconstitute ONM proteins such as nesprins.

### The C-termini of SUN1 and SUN2 inserted into ANMs remain solvent-exposed

Since the LINC complex assembly is driven by the direct interaction of the C-terminal SUN domain of SUN proteins with the C-terminal KASH peptide of KASH proteins within the PNS, we next needed to determine the orientation of EGFP-SUN1^FL^-His_6_ and EGFP-SUN2^FL^-His_6_ in reconstituted ANMs. To do this, we developed an imaging-based protease protection assay using the *Streptomyces griseus*-derived pronase (Xu et al., 1988), which was added to reconstituted ANMs on beads after 8 hours of incubation followed by a washing step. As EGFP is fused to the N-termini of our SUN protein constructs (Figure 3A), the loss of fluorescence following pronase addition would indicate that their N-termini are solvent-exposed. Alternatively, if the EGFP fluorescence were detected in the presence of pronase, we would conclude that the N-termini of our constructs was found within the space between the supported lipid bilayer of the SUPER template and the silica bead (Figure 3B).

**Figure 3:**
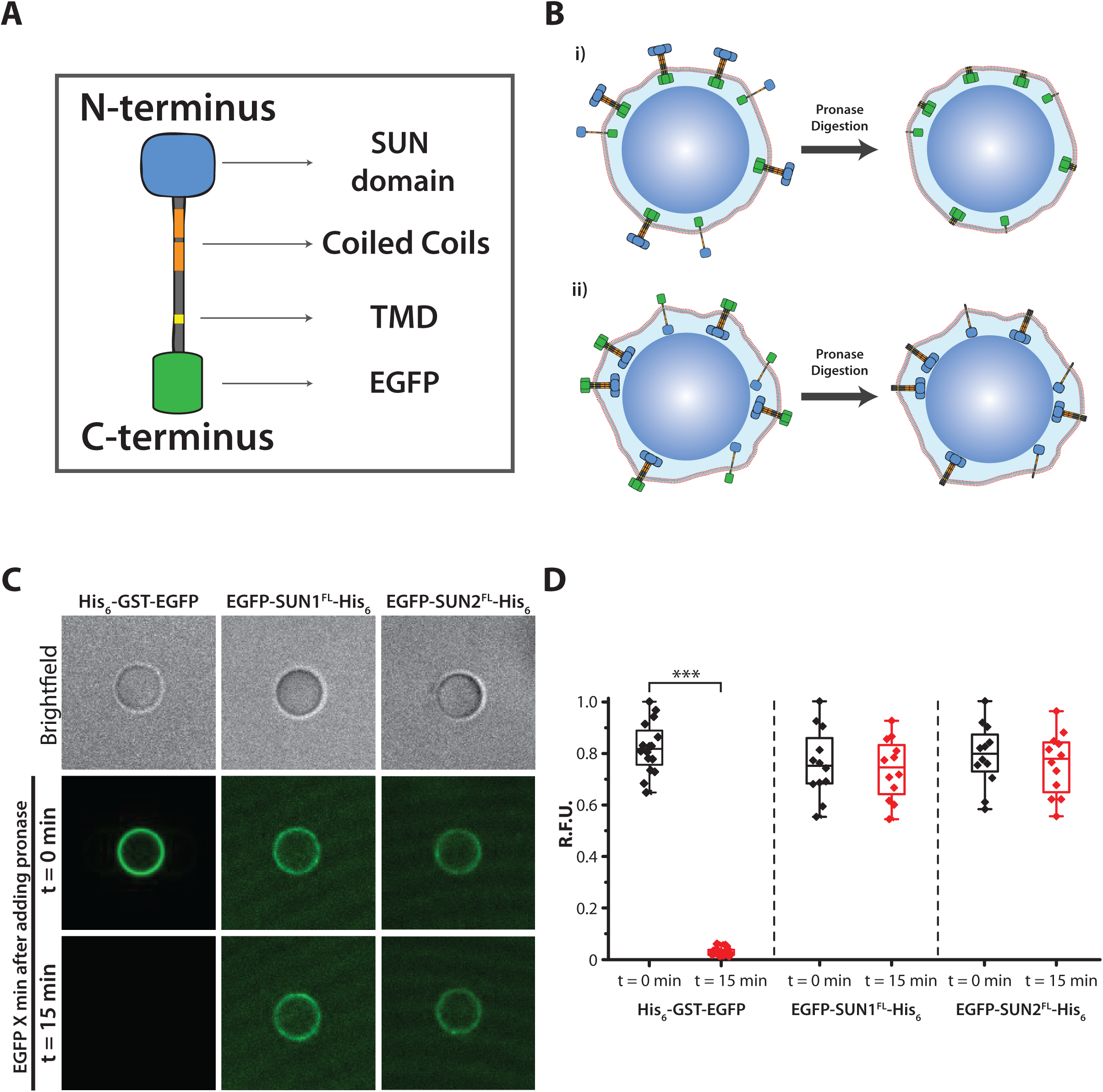
Orientation of CFE-synthesized FL SUN1 and SUN2 inserted in ANMs. **A)** Illustration of a FL SUN protein with EGFP fused to its N-terminus. **B)** Schematic of the pronase digestion assay used to determine the topology of ANM-inserted FL SUN proteins synthesized by CFE. If the SUN proteins were oriented with their N-termini protruding away from the ANM into the solution and their C-termini inserted in between the lipid bilayer and the silica bead, EGFP would be protected from pronase-mediated degradation (i). If SUN proteins were oriented in the opposite direction, EGFP would be degraded by pronase (ii). **C)** Representative images of SUPER templates incubated in CFE reactions expressing the indicated FL SUN protein constructs before (t = 0 min) or after (t = 15 min) the addition of pronase. As a positive control for pronase digestion, purified His_6_-GST-EGFP was incubated with SUPER templates containing 10% DOGS-NTA-Ni prior to the addition of pronase. Scale bar: 5 μm. **D)** Box plots depicting the relative fluorescence units (RFU) of EGFP quantified on ANMs before (t = 0 min) and after (t = 15 min) the addition of pronase for the indicated constructs from 3 independent experiments (n = 24 – 30 beads). *** *p* < 0.001.

To control for protease activity, we tested the ability of pronase to efficiently degrade His_6_-tagged glutathione-S-transferase (GST) EGFP (His_6_-GST-EGFP), the membrane-association of which was promoted by the incorporation of 10% 1,2-dioleoyl-*sn*-glycero-3-[(N-(5-amino-1-carboxypentyl)iminodiacetic acid)succinyl] (nickel salt) (DOGS-NTA-Ni) into the SUPER templates incubated in the CFE reaction (Figure 3B). As expected, the addition of pronase reduced the levels of EGFP fluorescence detected on the surface of His_6_-GST-EGFP-bound ANMs to negligible levels between 5-15 minutes (Figures 3C and S2). In contrast, no significant difference in EGFP fluorescence was detected on ANMs containing either EGFP-SUN1^FL^-His_6_ or EGFP-SUN2^FL^-His_6_ before or 15 minutes after the addition of pronase (Figures 3C-D). To further examine the orientation of EGFP-SUN1^FL^-His_6_ and EGFP-SUN2^FL^-His_6_ in ANMs, we performed the pronase protection assay on ANMs containing these reconstituted SUN proteins, which were labeled with an anti-His monoclonal antibody directly conjugated to the fluorescent dye Alexa Fluor 647 (Penta-His-AF647) (Fig. S3). Confocal images of these labeled SUN protein-containing ANMs prior to the addition of pronase revealed clear ANM-associated EGFP and Penta-His-AF647 fluorescence, while only EGFP fluorescence was detected on the ANMs after exposure to pronase. Thus, these results suggest that the N-termini of EGFP-SUN1^FL^-His_6_ and EGFP-SUN2^FL^-His_6_ reside within the space in between the supported lipid bilayer and the silica bead of the ANM.

### SUN1 contains three TMDs

Several lines of experimental evidence in the literature support a model where the SUN proteins are single-pass type II membrane proteins with nucleoplasmic N-termini and luminal C-termini (Sosa et al., 2013, Starr and Fridolfsson, 2010) However, earlier studies demonstrated the presence of additional hydrophobic regions (HRs) in the NDs of both SUN1 and SUN2 (Crisp et al., 2006, Hodzic et al., 2004, Liu et al., 2007, Turgay et al., 2010). To begin to assess whether or not additional TMDs exist in SUN1, we generated EGFP-and His_6_-tagged constructs encoding the amino acids N-and C-terminal of the previously identified TMD, which were referred to as the nucleoplasmic domain (ND) and luminal domain (LD), respectively (Figure 4A). Whereas CFE-synthesized EGFP-SUN1^LD^-His_6_ remained soluble, EGFP-SUN1^ND^-His_6_ strongly associated with SUPER templates incubated in CFE reactions (Figure 4B). These results clearly suggest that additional TMDs and/or HRs within the ND of SUN1.

**Figure 4:**
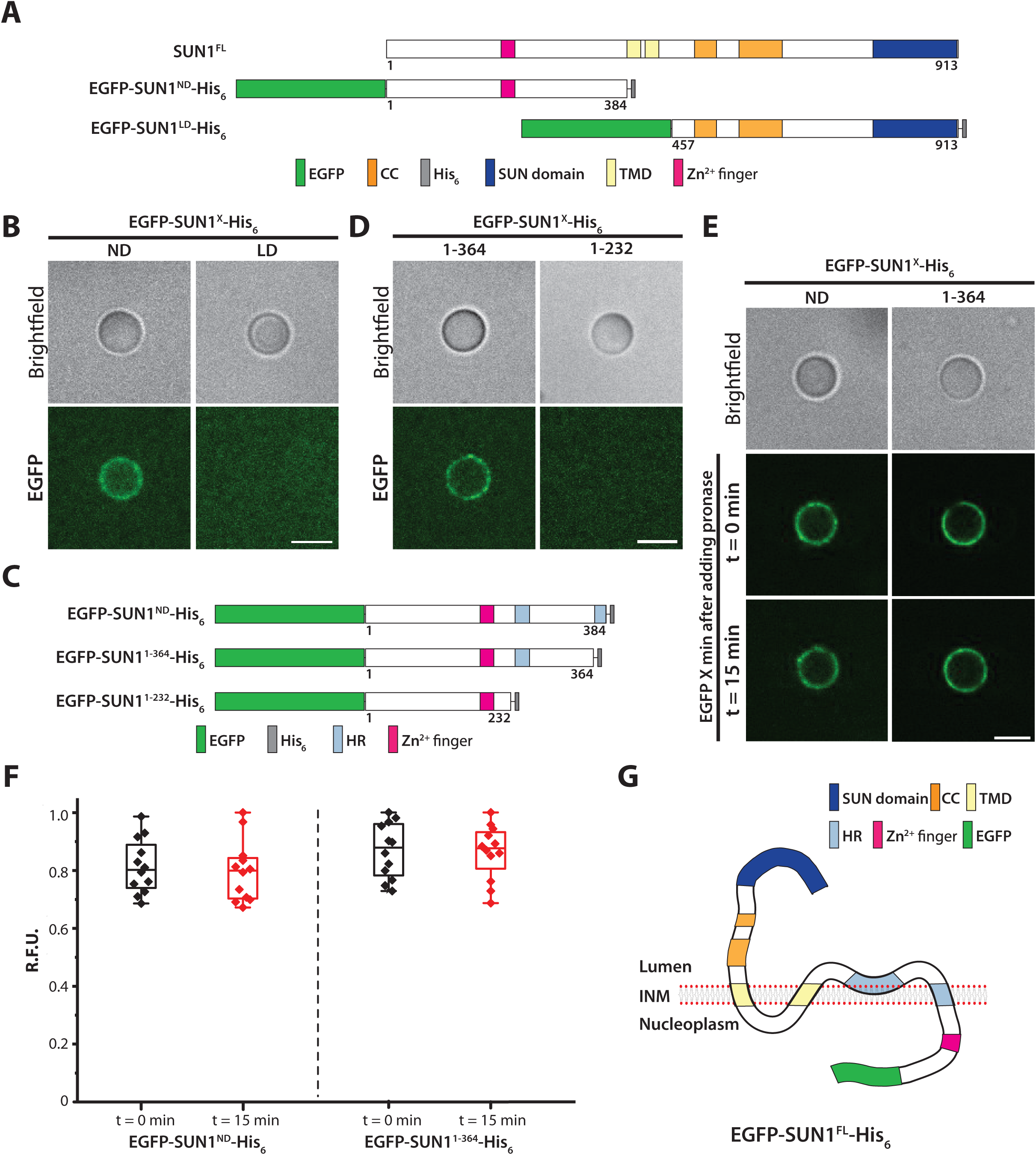
Topology of SUN1 inserted in ANMs. **A)** and **C)** Illustration of the constructs used in this figure. **B)**and **D)** Representative images of silica beads or SUPER templates incubated in CFE reactions of the indicated constructs. **E)** Representative images of SUPER templates incubated in CFE reactions expressing the indicated constructs before (t = 0 min) or after (t = 15 min) the addition of pronase. **F)** Box plots depicting the RFU of EGFP quantified on ANMs before (t = 0 min) and after (t = 15 min) the addition of pronase for the indicated constructs from 3 independent experiments (n = 12 beads per condition). **G)** Working model for the topology of EGFP-SUN1^FL^-His_6_ inserted into the INM. Scale bars in B, D, and E: 5 μm.

Computational analysis of the solvent accessibilities of the amino acid residues present in the ND of SUN1 using the SABLE server (http://sable.cchmc.org/sable_doc.html) (Adamczak et al., 2004) revealed the presence of two additional HRs in the SUN1^ND^ (Figure 4C). To test the role of these HRs in promoting the association of SUN1^ND^ with ANMs, we generated a panel of EGFP-and His_6_-tagged constructs encoding SUN1^ND^ truncations (Figure 4C). The single HR-containing EGFP-SUN1^1-364^-His_6_ construct still associated with ANMs, whereas the HR-devoid EGFP-SUN1^1-232^-His_6_ construct did not (Figure 4D). Based on these results, we conclude that the HRs present in SUN1^ND^ mediates its ability to associate with the ANM. While the SUN1^ND^ also contains an enigmatic C2H2 Zn^2+^ finger (Liu et al., 2007, Padmakumar et al., 2005), it is not sufficient for ANM association.

To determine if the HRs found within SUN1^ND^ are TMDs or peripherally membrane-associated, we again turned to the pronase protection assay described above. Treatment of EGFP-SUN1^ND^-His_6_ or EGFP-SUN1^1-364^-His_6_-associated ANMs with pronase for 15 minutes did not significantly decrease the levels of ANM-associated EGFP fluorescence (Figures 4E-F). In addition, we performed pronase protection assays on ANMs containing either EGFP-SUN1^ND^-His_6_ or EGFP-SUN1^1-364^-His_6_, which were also labeled with Penta-His-AF647 (Fig. S3). Similar to what was observed with ANMs containing either EGFP-SUN1^FL^-His_6_ or EGFP-SUN1^FL^-His_6_, confocal images of these labeled SUN protein-containing ANMs prior to the addition of pronase revealed clear ANM-associated EGFP and Penta-His-AF647 fluorescence, while only EGFP fluorescence was detected on the ANMs after exposure to pronase. Thus, we propose that one of the two additional HRs in the SUN1^ND^ is a TMD. Based on these results and the possibility of a membrane-associating HR in SUN1^ND^ a potential model of SUN1 topology in the INM is shown (Figure 4G) which would preserve the luminal orientation of the C-terminus while maintaining the nucleoplasmic orientation of the N-terminus.

### SUN2 contains a single TMD and a membrane-associated HR

We next asked if SUN2 also possessed more than one TMD by testing the ability of EGFP-and His_6_-tagged constructs encoding the SUN2^ND^ and SUN2^LD^ to associate with membranes (Figure 5A). Similar to SUN1, EGFP-SUN2^LD^-His_6_ remained soluble, while EGFP-SUN2^ND^-His_6_ strongly associated with SUPER templates incubated in CFE reactions (Figure 5B). Computational analysis using the SABLE server revealed the presence of a single HR within the ND of SUN2 (Figure 5C), which was found to be critical for SUN2^ND^ to associate with membranes due to the fact that a construct lacking this HR, EGFP-SUN2^1-131^-His_6_, remained soluble (Figure 5D). However, the addition of pronase to EGFP-SUN2^ND^-His_6_-associated SUPER templates reduced EGFP fluorescence to background levels (Figure 5E-F). Consequently, we propose that the SUN2^ND^ peripherally associates with the nucleoplasmic leaflet of the INM, thereby maintaining the type II membrane protein topology of SUN2 (Figure 5G).

**Figure 5:**
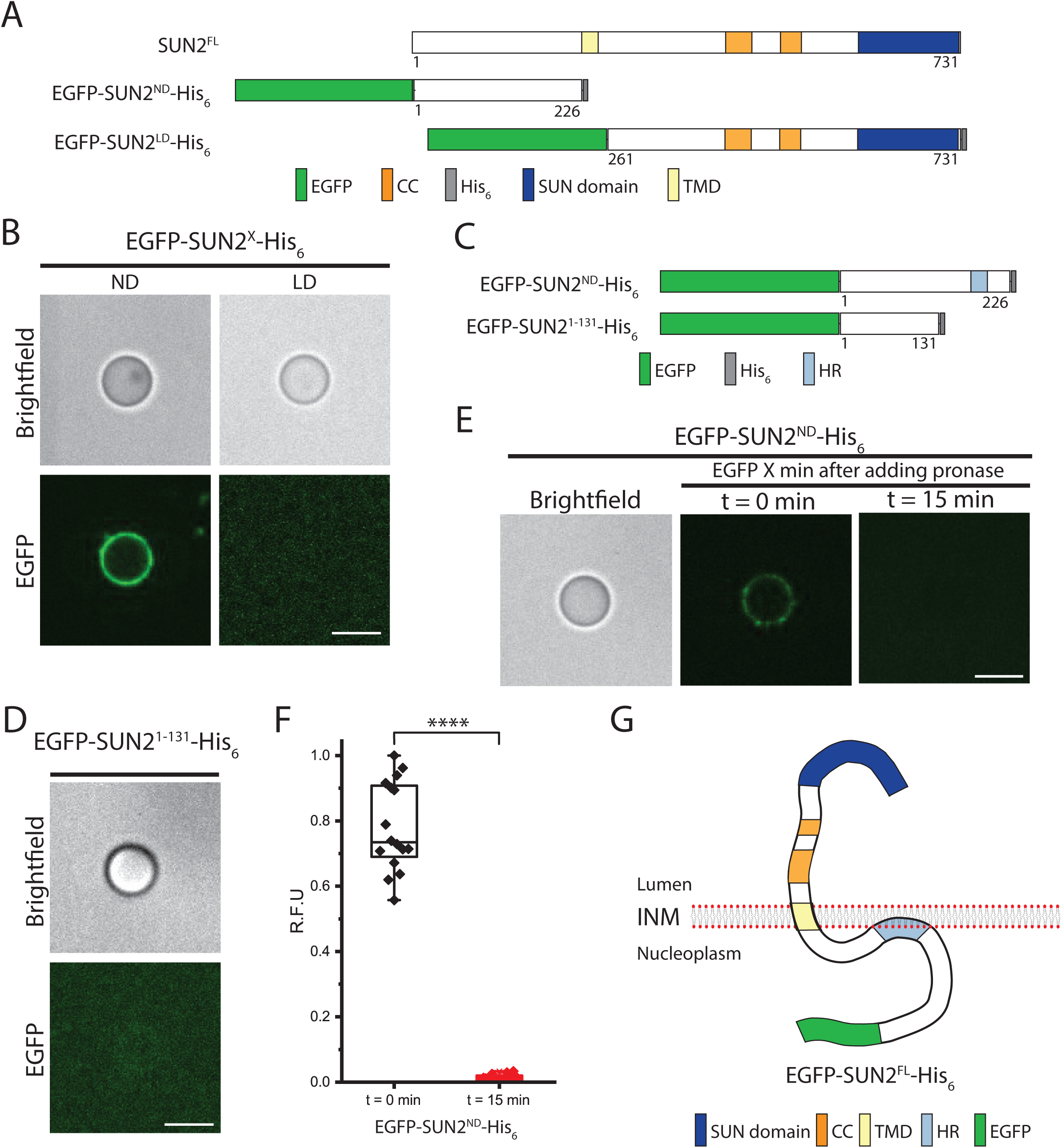
Topology of SUN2 inserted in ANMs. **A)** and **C)** Illustration of the constructs used in this figure. **B)** and **D)** Representative images of SUPER templates incubated in CFE reactions of the indicated constructs. **E)** Representative images of SUPER templates incubated in CFE reactions expressing the indicated constructs before (t = 0 min) or after (t = 15 min) the addition of pronase. **F)** Box plots depicting the RFU of EGFP quantified on ANMs before (t = 0 min) and after (t = 15 min) the addition of pronase for the indicated constructs from 3 independent experiments (n = 12 beads per condition). *** *p* < 0.001. **G)** Working model for the topology EGFP-SUN2^FL^-His_6_ inserted into the INM. Scale bars in B, D, and E: 5 μm.

### Reconstitution of KASH-binding FL SUN1 and SUN2 in ANMs

Having demonstrated that we could successfully use CFE to generate supported artificial lipid bilayers containing FL SUN1 and SUN2 oriented such that their C-termini were solvent-exposed, we finally ask whether or not we could use these ANMs to reconstitute LINC complex assembly *in vitro*. Since the core of the LINC complex is formed by the direct interaction between the LD of SUN proteins and the KASH peptide of KASH proteins (Sosa et al., 2013) (Figure 6A), we tested the ability of EGFP-SUN1^FL^-His_6_-and EGFP-SUN2^FL^-His_6_-containing ANMs to recruit wild-type (WT) KASH peptides from the KASH protein nesprin-2 with tetramethylrhodamine (TRITC) attached to their N-termini (TRITC-KASH2^WT^). ANMs containing EGFP-SUN1^FL^-His_6_ or EGFP-SUN2^FL^-His_6_ displayed an efficient recruitment of TRITC-KASH2^WT^. Importantly, neither ANM was able to recruit TRITC-labeled KASH peptides lacking the four C-terminal amino acids that are required for the SUN-KASH interaction to occur normally (Padmakumar et al., 2005) (TRITC-KASH2^ΔPPPT^) to levels similar to those observed with TRITC-KASH2^WT^ (Figures 6B-C). However, the recruitment of TRITC-KASH2^ΔPPPT^ to ANMs containing EGFP-SUN1^FL^-His_6_ or EGFP-SUN2^FL^-His_6_ was slightly elevated relative to its recruitment to SUPER templates, which were not incubated in CFE reactions. Therefore, TRITC-KASH2^ΔPPPT^ appears to maintain some ability to interact with FL SUN proteins. Because LINC complex assembly is critically dependent upon the ability of SUN proteins to directly interact with KASH peptides, these results strongly suggest that we have successfully reconstituted the SUN-KASH interaction using CFE and ANMs.

**Figure 6:**
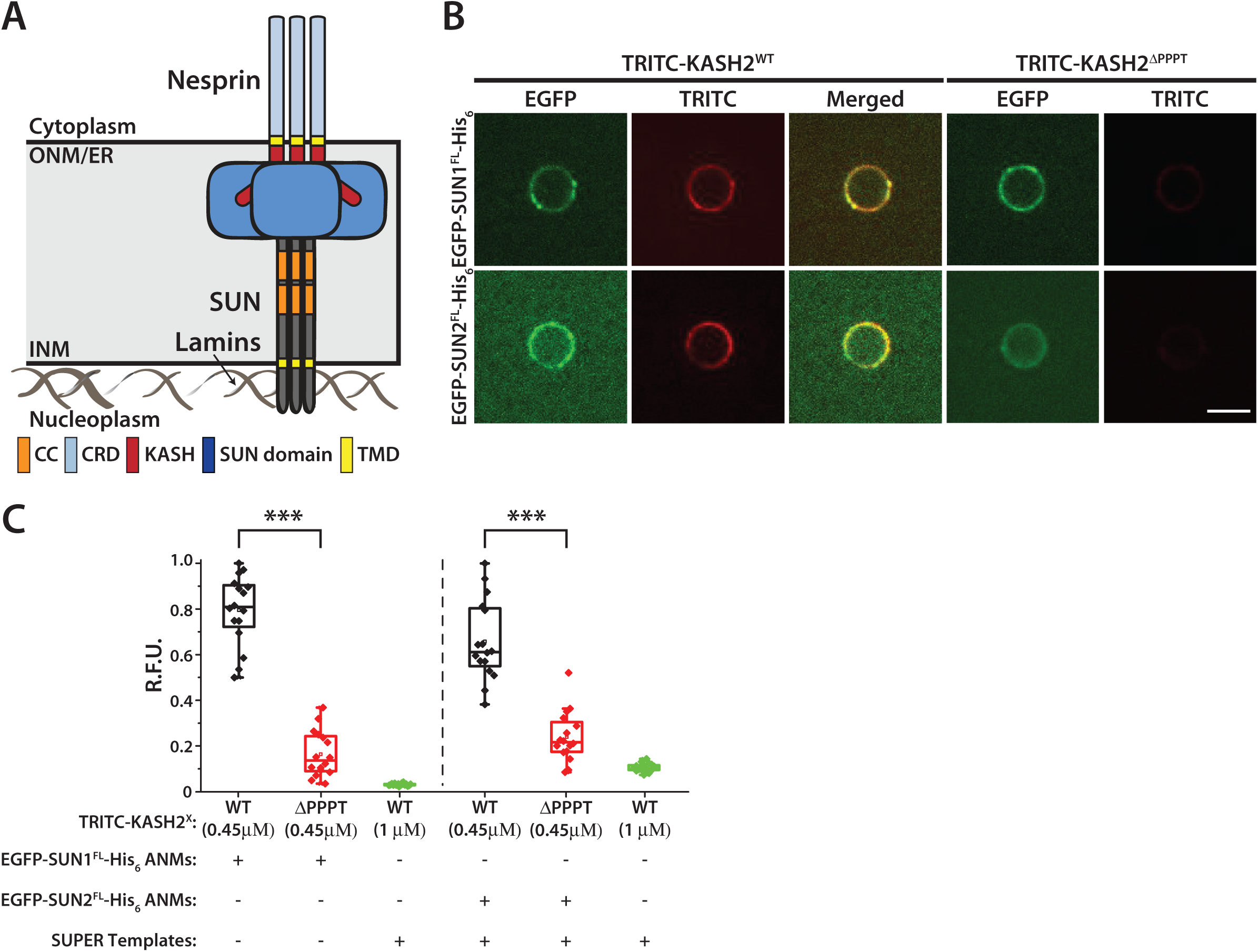
Reconstitution of KASH-binding SUN1^FL^ and SUN2^FL^ complexes using CFE and ANMs. **A)** Schematic depicting an assembled LINC complex based on previously published structural and computational modeling studies (Jahed et al., 2018, Sosa et al., 2012). CRD: central rod domain. **B)** Representative images - of ANMs containing the indicated inserted SUN protein as well as SUPER templates incubated together with the indicated KASH2 peptide. Scale bar: 5 μm. **C)** Box plots depicting the quantified RFUs from the indicated KASH2 peptides recruited to ANMs containing the inserted indicated SUN protein as well as to SUPER templates, which were not incubated in CFE reactions from 3 independent experiments (n = 12 beads per condition). Scale bar for B): 5 μm. *** *p* < 0.001.

## DISCUSSION

Here, we describe ANMs as a bottom-up synthetic biology platform for the reconstitution and mechanistic dissection of LINC complex assembly. We show that we can reconstitute FL SUN1 and SUN2 proteins into ANMs with their N-terminal NDs oriented in between the lipid bilayer and the silica bead while their C-terminal LDs extend away from the ANM and into the solution. We also use ANMs to determine that SUN2 possesses a single transmembrane domain, while SUN1 possesses three. Finally, we demonstrate that ANM inserted FL SUN proteins are capable of recruiting the KASH peptide of nesprin-2, suggesting that we have successfully reconstituted the SUN-KASH interaction, which is the core of the LINC complex.

The minimal ability of TRITC-KASH2^ΔPPPT^ to interact with FL SUN proteins inserted into ANMs suggests that the PPPT motif is not the only binding interface between the KASH2 peptide and SUN proteins. In fact, KASH2 interacts with SUN2 through two additional binding interfaces (Sosa et al., 2012 *Cell* and Cain et al., 2018 *Curr Biol*). The first interface consists of residues -4 to -14 of the KASH peptide, which insert into a groove formed between two SUN2 protomers with the conserved hydrophobic residues at -7 and -9 being oriented towards the groove. The third interface, residues -15 to -23, fit along the surface of the adjacent SUN2 protomer. A conserved cysteine at position -23 of the KASH peptide is oriented to form an intermolecular disulfide bond with a conserved cysteine in the SUN domain of SUN2. Future experiments aimed at testing the role of these other binding interfaces on the residual recruitment of TRITC-KASH2^ΔPPPT^ to reconstituted SUN protein-containing ANMs will help address this question.

The orientation of SUN proteins reconstituted in ANMs is not random, presumably due to the fact that the CFE reaction lysates contain ER-derived microsomes, which contain the cellular machinery necessary for the co-translational translocation of membrane proteins (Brödel et al., 2015). Since the interior of ER-derived microsomes is equivalent to the ER lumen and the contiguous perinuclear space of the nuclear envelope, the C-terminal LD of SUN proteins will be co-translationally translocated into the interior while the N-terminal ND will remain on the outside of the ER-derived microsomes in the CFE reaction. Thus, when the SUPER templates are introduced into the SUN protein-expressing CFE reactions, the SUN protein-containing ER-derived microsomes fuse with the supported lipid bilayer surrounding the silica bead of the SUPER template. Consequently, the N-terminal ND can now be found in the space between the lipid bilayer and the silica bead, while the C-terminal LD is oriented away from the ANM and solvent-exposed. However, this case does not hold for EGFP-SUN2^ND^, which contains a membrane-associated HR, due to the fact that the protein is likely synthesized outside of the ER-derived microsomes and is only present on the solvent-exposed side of the ANM.

Our finding that SUN1 possesses three TMDs stands in discord with a previous report, which concluded that SUN1 is a type II membrane protein with a single TMD based on in situ proteinase K digestions performed in HeLa cells expressing mouse SUN1 tagged with HA at their N-termini and EGFP followed by a myc epitope at their C-termini, which were treated with low concentrations of digitonin (Crisp et al., 2006, Liu et al., 2007, Padmakumar et al., 2005). The effect of proteinase K was subsequently assessed by Western blot analysis following myc immunoprecipitation as well as by immunofluorescence microscopy. Even though two of the SUN1 HRs could not be ruled out as being membrane-associated in these experiments, the authors concluded that SUN1 was a type II membrane protein with a single TMD. Since EGFP-SUN1^1-364^-His_6_ contains this HR and retains EGFP fluorescence in the presence of pronase, our results suggest that the first HR is in fact a TMD. We propose that the second HR peripherally associates with the luminal leaflet of the INM, where it may serve as a docking-site for an as-of-yet unidentified SUN1-binding partner present within the perinuclear space of the NE. The presence of multiple TMDs in SUN1 and not in SUN2 may be relevant during meiosis, when SUN1-containing LINC complexes associate with the telomeres of meiotic chromosomes to mediate their microtubule-dependent movement during homologue pairing via cytoplasmic dynein (Ding et al., 2007, Horn et al., 2013, Morimoto et al., 2012). The additional TMDs may help buffer the shear and tensile forces applied to telomere-associated SUN1-containing LINC complexes by dyneins, which walk along microtubules oriented parallel to the nuclear envelope (Jahed and Mofrad, 2018). It will be interesting to determine if the additional TMDs in SUN1 are required for this process.

The physiological relevance of the ability of the ND of SUN2 to peripherally associate with the nucleoplasmic leaflet of the INM remains unknown. One intriguing possibility is that the peripheral membrane-association of SUN2 may regulate its homo-oligomerization, as has been observed with HIV-1 Gag and matrix proteins (Wegener and Campbell, 2008) as well as Ras GTPases (Chavan et al., 2015). Another may be related to the targeting of SUN proteins to the INM following their synthesis in the ER/ONM. In order to gain access to the INM, newly synthesized SUN proteins utilize nuclear targeting sequences to move across NPCs through a poorly defined mechanism (Tapley et al., 2011, Turgay et al., 2010). Recently, the size of the ND of SUN proteins was estimated to be less than 10 nm in diameter (Jahed et al., 2016), which is near the upper size limit for the transport of the NDs of INM proteins through the NPC (Lin et al., 2016, Ungricht and Kutay, 2015). Thus, the additional TMDs and HR identified in SUN1 and SUN2 may serve to reduce the footprint of their NDs so that they can properly target the INM.

It is worth noting that unlike His_6_-GST-EGFP, which evenly coats the surface of the SUPER templates, ANM-inserted EGFP-SUN1^FL^-His_6_ and EGFP-SUN2^FL^-His_6_ appear to form clusters within the membranes. These clusters may represent higher-order SUN protein oligomers such as those previously suggested for SUN1 (Hennen et al., 2018, Jahed et al., 2018, Lu et al., 2008). Alternatively, these SUN protein clusters may highlight the existence of membrane nanodomains within the ANMs (Krapf, 2018), which can be explored by altering the composition of lipids used to generate the SUPER templates. Moreover, the ND SUN constructs appear to be able to form fluorescent patches in ANMs similar to the FL SUN proteins. However, the ability of EGFP-SUN1^ND^ and EGFP-SUN2^ND^ to form these patches is not necessarily mutually exclusive with the previously proposed mechanisms of LD-mediated higher-order SUN protein clustering (Hennen et al., 2018, Jahed et al., 2018, Wang et al., 2012, Zhou et al., 2012). Based on our results, we suggest that both the LDs and NDs of SUN proteins can separately homo-oligomerize within the perinuclear space of the nuclear envelope and nucleoplasm, respectively. It will be interesting to test this hypothesis experimentally in future studies.

### Future directions

ANMs will allow for testing proposed mechanisms for regulating LINC complex assembly and/or disassembly in cells, which have been difficult to address experimentally. For example, the modulation of the oligomeric states of the CC-containing regions present within the LDs of SUN proteins was recently proposed as a potential target for the regulation of LINC complex assembly (Nie et al., 2016), as the homo-oligomerization of the SUN2 LD is essential for KASH peptide-binding (Sosa et al., 2012). Since these studies were performed using purified fragments of the LD of SUN2, it is essential that they be repeated using FL SUN proteins inserted into ANMs. Alterations in the levels of intracellular calcium also represent a potential regulator of LINC complex assembly, as the recently identified cation loop present within the SUN domain is required for the SUN-KASH interaction (Sosa et al., 2012). While deciphering the effects of altered intracellular calcium levels on the engagement of SUN and KASH proteins within the PNS remains a significant technical challenge, the solvent exposed LDs of FL SUN1 and SUN2 on the surface of ANMs provides a simplified system for testing the effect of manipulating calcium levels on LINC complex assembly.

In addition, ANMs may help quantitatively assess the ability of KASH peptides from different nesprins to interact with SUN proteins. This will be especially interesting for the KASH peptides of KASH5 and lymphoid-restricted membrane protein, which unlike the KASH peptides of nesprins 1-3 lack the intermolecular SUN-KASH disulfide bond forming conserved cysteine present at position -23 (Horn et al., 2013). Despite being non-essential for the SUN-KASH interaction, we recently showed that the conserved cysteines present in SUN and nesprin proteins are required for actin-dependent nuclear anchorage, yet dispensable for microtubule-dependent nuclear movement (Cain et al., 2018). Therefore, KASH peptides containing this conserved cysteine may display differential SUN protein-binding kinetics compared to those that do not. Furthermore, ANMs may finally provide the long sought-after experimental system for defining the mechanism of LINC complex regulation by the luminal ATPase torsinA (Saunders et al., 2017, Saunders and Luxton, 2016), due to the lack of a need for protein purification.

Besides SUN1 and SUN2, mammals express three additional testis-specific SUN proteins (SUN3, SUN4, and SUN5) (Sosa et al., 2013). While immunoprecipitation experiments show that SUN3 can associate with nesprin-1 and nesprin-3 (Göb et al., 2010), the ability of SUN4 and SUN5 to interact with KASH proteins remains unknown. Nevertheless, ANMs provide an ideal experimental platform for future explorations of the ability of these poorly studied SUN proteins to form LINC complexes.

The use of ANMs is by no means limited to SUN and KASH proteins. ANMs can provide a simple and powerful experimental platform for structure/function analyses of any membrane protein, nuclear or otherwise. By bridging the gap between *in vitro* studies using purified proteins and *in vivo* imaging-based biochemical studies, ANMs open the door to a previously inaccessible set of experiments designed to provide mechanistic insights into protein behavior, which has the potential to become invaluable in this post-genomic age. Akin to other attempts to use modular approaches for building artificial cells or reconstituting cellular processes (Liu and Fletcher, 2009, Majumder and Liu, 2017), future efforts will be aimed at reconstituting the nuclear lamina and its interaction with the nucleoplasmic domain of SUN proteins within an ANM. Finally, it might be possible to use ANMs as templates for the reconstitution of an artificial nuclear envelope through a “layer-by-layer” double membrane assembly approach using poly L-lysine as an electrostatic polymer linker, which was previously shown to form lipid multilayers via vesicle rupture onto existing supported lipid bilayers (Heath et al., 2016). Due to the ability of the nesprin KASH peptide to directly interact with the SUN protein SUN domain (Ostlund et al., 2009, Sosa et al., 2012), ruptured nesprin-containing ER-derived microsomes generated by CFE would presumably coat the ANM containing the reconstituted SUN protein such that the nesprin N-terminus would extend into the solution while its C-terminus would face the ANM.

## ACKNOWLEDGMENTS

This work was financially supported in part by the Dystonia Medical Research Foundation (GWGL) and the NSF (APL (MCB-1612917)). MSD was supported by the NSF Graduate Research Fellowship Program. We also thank Nick Emery for assisting with the purification of His_6_-GST-EGFP.

## ABBREVIATIONS

ANM: Artificial nuclear membrane
CC: Coiled-coil
CFE: Cell-free expression
CRD: Central rod domain
DiI: 1,1’-Dioctadecyl-3,3,3’,3’-Tetramethylindocarbocyanine Perchlorate
DOGS-NTA-Ni: 1,2-dioleoyl-*sn*-glycero-3-[(N-(5-amino-1-carboxypentyl)iminodiacetic acid)succinyl] (nickel salt)
DOPC: 1,2-dioleoyl-*sn*-glycero-3-phosphatidylcholine
DOPE: 1,2-dioleoyl-*sn*-glycero-3-phosphoethanolamine
DOPS: 1,2-dioleoyl-*sn*-glycero-3-phospho-L-serine (sodium salt)
Egg-PA: L-α-phosphatidic acid (Egg, Chicken) (sodium salt)
ER: Endoplasmic reticulum
F: Forward
FL: Full-length
KASH: Klarsicht, ANC-1, SYNE homology
HR: Hydrophobic region
INM: Inner nuclear membrane
KLD: Kinase, Ligase, *Dpn*I
LD: Luminal domain
LINC: Linker of nucleoskeleton and cytoskeleton
NE: Nuclear envelope
ND: Nucleoplasmic domain
NPC: Nuclear pore complex
ONM: Outer nuclear membrane
PNS: Perinuclear space
R: Reverse
RE: Restriction enzyme
RFU: Relative fluorescence units
SUN: Sad1/UNC-84
SUPER: Supported bilayers with excess membrane reservoir
SUV: Small unilamellar vesicle
TMD: Transmembrane domain
TRITC: Tetramethylrhodamine
WT: Wild-type

## MATERIALS AND METHODS

### Reagents

Restriction enzymes were either purchased from New England Biolabs (NEB, Ipswich, MA) or Promega (Madison, WI). Phusion DNA polymerase, T4 DNA ligase, and PNK were also purchased from NEB. All other chemicals were from Sigma-Aldrich (St. Louis, MI) unless otherwise specified. Wizard SV Gel and PCR Clean-Up System was from Promega. All lipids were purchased from Avanti Polar Lipids Inc. (Alabaster, AL) in chloroform stock solutions. Anti-EGFP antibody (ab6556) was purchased from Abcam (Cambridge, MA) and used at 1:500 for Western blotting. DiI was purchased from Thermo Fisher Scientific (Waltham, MA) and used at 1:20. Penta-His-AF647 was purchased from QIAGEN (Hilden, Germany) and used at 1:60 for immunofluorescence.

### Cell culture

HeLa S3 cells obtained from the ATCC (Manassas, VA) were cultured using standard sterile technique in SMEM medium supplemented with 5% bovine calf serum (Sigma-Aldrich), 0.5% penicillin/streptomycin, 10 mM HEPES (pH 7.4) and 0.5% glutamax (Thermo Fischer Scientific, Waltham, MA).

### CFE lysate generation

HeLa S3 cells were cultured in a flat bottom bell jar with rotating flaps using a magnetic stirrer until reaching a density of ~6 × 10^6^ cells/ml. Harvested cells were washed three times with washing buffer (35 mM HEPES pH 7.5, 140 mM NaCl and 11 mM glucose) and once with extraction buffer (20 mM HEPES pH 7.5, 45 mM potassium acetate, 45 mM potassium chloride, 1.8 mM magnesium acetate, 1 mM DTT). The washed cells were then resuspended in 1 ml of extraction buffer before being lysed in a BeadBug Homogenizer (Benchmark Scientific, Edison, NJ) using 2 ml tubes filled partially with 0.1 mm titanium beads. To remove debris, nuclei, and most organelles, the resulting lysate was centrifuged three times at 16,000 xg, after which 50 μl aliquots of the cleared lysate were flash-frozen in liquid nitrogen and stored at -80°C for future use.

### CFE reactions

CFE reactions were performed by mixing 9 μl of the CFE lysate, 2.25 μl Mix 1 (27.6 mM magnesium acetate, 168 mM HEPES (pH 7.5)) and 2.7 μl GADD34 (final concentration of 310 nM) in a 1.5 mL microcentrifuge tube, which was then incubated at 32°C for 10 minutes. Thereafter, 2.25 μl Mix 2 (12.5 mM ATP, 8.36 mM GTP, 8.36 mM CTP, 8.36 mM UTP, 200 mM creatine phosphate, 7.8 mM HEPES (pH 7.5), 0.6 mg/ml creatine kinase, 0.3 mM amino acid mixture, 5 mM spermidine, and 44.4 mM DTT), 5 nM of plasmid DNA, and 1.8 μl T7 RNA polymerase (final concentration of 450 nM) were added to the CFE reaction followed by vortexing. All supplements for the CFE reaction were stored at -80°C for future use.

### ANM generation

SUPER templates were generated as previously described (Neumann et al., 2013). Briefly, SUVs composed of 45% 1,2-dioleoyl-*sn*-glycero-3-phosphatidylcholine (DOPC), 27% 1,2-dioleoyl-*sn*-glycero-3-phosphoethanolamine (DOPE), 9% 1,2-dioleoyl-*sn*-glycero-3-phospho-L-serine (sodium salt) (DOPS), 2.2% L-α-phosphatidic acid (Egg, Chicken) (sodium salt) (Egg-PA), and 16.8% cholesterol were prepared by extrusion in milli-Q water through a 100 nm extruder (T&T Scientific, Knoxville, TN) and then fused with 5 μm silica beads (Bangs Laboratories, Fischers, IN) in the presence of 1M NaCl. The resulting SUPER template beads were washed with milli-Q water twice and then resuspended in 30 μl of milli-Q water at a final concentration of ~ 9.6 × 10^6^ beads/ml, 2 μl of which were then added to a CFE reaction and incubated for 8 hours at 32°C.

### DNA constructs

The His_6_-GST-EGFP pET28 construct was a kind gift from Michael Jewet (Northwestern University, Evanston, IL). A previously described EGFP-tagged mouse SUN1^FL^ construct (EGFP-SUN1^FL^) (Luxton et al., 2010, Ostlund et al., 2009) was used as a template for the generation of the SUN1 constructs used in this study. Initially, SUN1^FL^ was PCR amplified from EGFP-SUN1^FL^ using the primers SUN1^FL^-Forward(F) and SUN1^FL^-Reverse (R), which contain 5’ *Eco*RI and *Sal*I cut sites, respectively. The PCR product was purified and digested alongside pT7CFE1-Chis (ThermoFischer Scientific) with *Eco*RI and *Sal*I. Following gel purification, the digested PCR product and plasmid were ligated together to create SUN1^FL^-His_6_. Unfortunately, an unwanted stop codon was found between SUN1^FL^ and the His_6_ tag. We removed this stop codon by PCR using the primers SUN1^FL/ΔSTOP^-F and SUN1^FL/ΔSTOP^-R and subsequent Kinase, Ligase, *Dpn*I (KLD) treatment where 2μL of the PCR product was treated with T4 ligase, T4 PolyNucleotide Kinase (PNK), and *Dpn*I in T4 ligase buffer in a 20μL reaction for 20 minutes at room temperature. To generate EGFP-SUN1^FL^-His_6_, EGFP was PCR amplified from EGFP-SUN1^FL^ using the primers EGFP-F and EGFP-R, both of which contain 5’ *Eco*RI cut sites. Following digestion with *Eco*RI, both the PCR product and SUN1^FL^-His_6_ were gel purified and ligated together to create EGFP-SUN1^FL^-His_6_. EGFP-SUN1^ND^-His_6_ was made by PCR amplifying the SUN1^ND^ using the primers EGFP-SUN1^ND^-F and EGFP-SUN1^ND^-F, which contain 5’ *Hind*III and *Kpn*I cut sites, respectively. The resulting PCR product was purified and digested alongside pT7CFE1-Chis with *Hind*III and *Kpn*I, both of which were gel purified, and ligated together to create EGFP-SUN1^ND^-His_6_. EGFP-SUN1^LD^-His_6_ was generated by PCR amplification using the primers EGFP-SUN1^LD^-His_6_-F and EGFP-SUN1^LD^-His_6_-R followed by purification and KLD treatment. EGFP-SUN1^1-364^-His_6_ and EGFP-SUN1^1-232^-His_6_ were generated similarly. While the same F primer, SUN1^FL/ΔSTOP^-F, was used for both constructs, the R primers used to create EGFP-SUN1^1-364^-His_6_ and EGFP-SUN1^1-232^-His_6_ were EGFP-SUN1^1-364^-His_6_-R and EGFP-SUN1^1-364^-His_6_-R, respectively.

A previously described EGFP-tagged mouse SUN2^FL^ construct (EGFP-SUN2^FL^) (Luxton et al., 2010, Ostlund et al., 2009) was used as a template for the generation of the SUN2 constructs used in this study. EGFP-SUN2^FL^-His_6_ was created in an analogous manner to EGFP-SUN1^FL^-His_6_. We first made SUN2_FL_-His6 using the primers SUN2^FL^-F and SUN2^FL^-R, which respectively contain 5’ *Eco*RI and *Xho*I cut sites, to PCR amplify SUN2^FL^ from EGFP-SUN2^FL^. The resulting PCR product and pT7CFE-Chis were both digested with *Eco*RI and *Xho*I, subsequently gel purified, and then ligated together to create SUN2^FL^-His_6_. Again, an unwanted stop codon was found between SUN2^FL^ and the His_6_ tag, which was removed by PCR using the primers SUN2^FL/ΔSTOP^-F and SUN2^FL/ΔSTOP^-R and subsequent KLD treatment. As described above, EGFP was PCR amplified from EGFP-SUN1^FL^ using the primers EGFP-F and EGFP-R and *Eco*RI digested alongside SUN2^FL^-His_6_, both of which were gel purified and ligated together to form EGFP-SUN2^FL^-His_6_. EGFP-SUN2^ND^-His_6_ and EGFP-SUN2^LD^-His_6_ were made by PCR amplification using the respective primer pairs EGFP-SUN2^ND^-His_6_-F/EGFP-SUN2^ND^-His_6_-R and EGFP-SUN2^LD^-His_6_-F/EGFP-SUN2^LD^-His_6_-R followed by purification and KLD treatment. EGFP-SUN2^1-131^-His_6_ was made by PCR amplification using the primers SUN2^FL/ΔSTOP^-F and EGFP-SUN2^1-131^-His_6_-R, the product of which was purified and subjected to KLD treatment. All of the above-mentioned primers are described in Table 1.

**Table 1:**
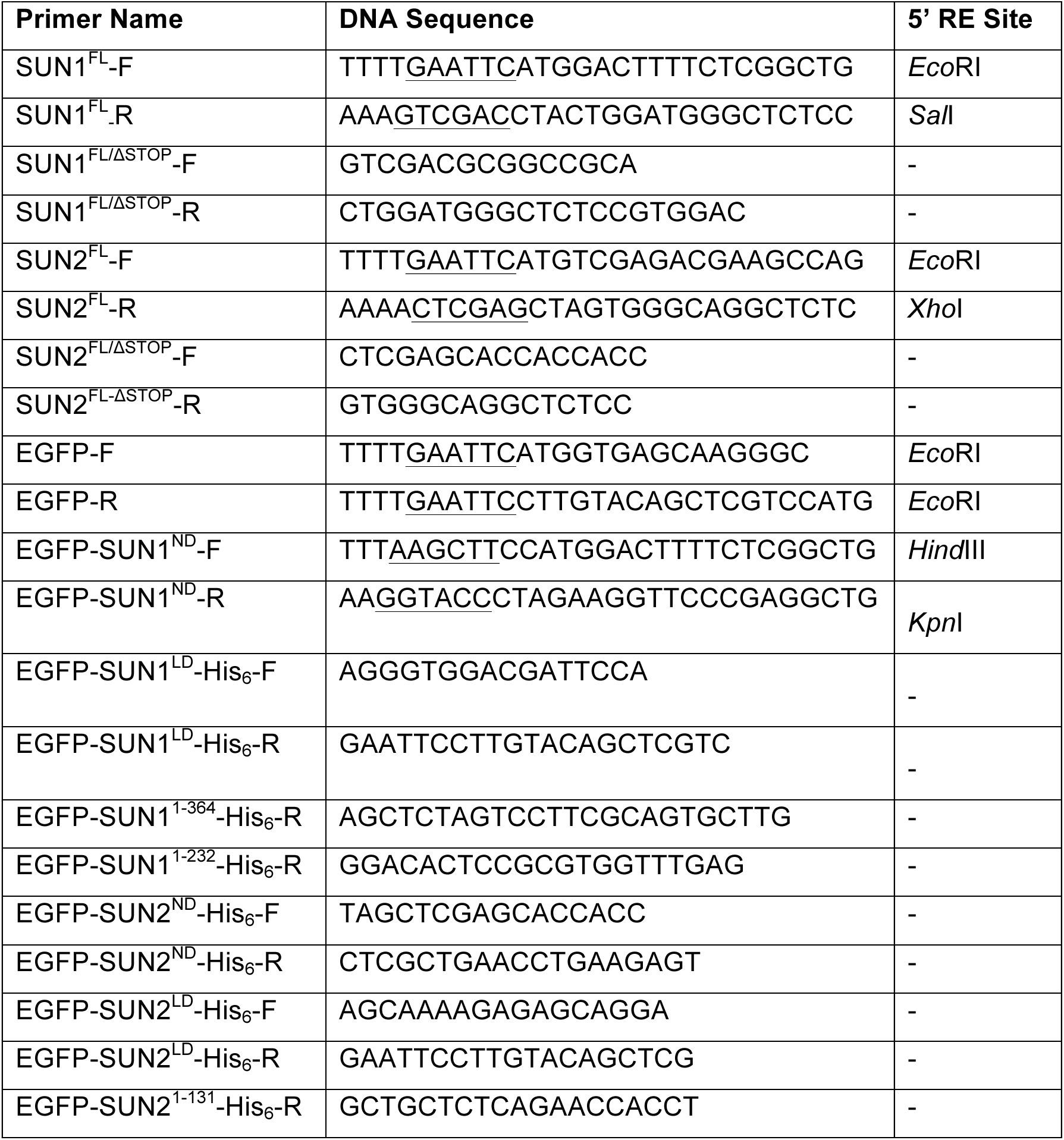
Primers used to generate the constructs used in this paper. The F or R in the primer name refers to forward or reverse, respectively. Restriction enzyme (RE) cut sites are underlined. The sequence encoding the linker is bolded.

### Protein purification

GST-tagged T7 RNA polymerase and truncated GADD34 were expressed in BL21 DE3 strain and purified using standard glutathione-based affinity chromatography as previously described (Mikami et al., 2008, Mikami et al., 2010). The His6-GST-EGFP construct was transformed into BL21 DE3 *E. coli* strain. 1 L liquid culture was inoculated with a single colony and induced with IPTG at OD600. Following 4 hours of growth at 37°C, cells were harvested at 10,000 xg for 15 minutes. The resulting cell pellet was then resuspended in 40 mL lysis buffer (20 mM HEPES (pH 7.5), 200 mM NaCl, 1 mM DTT, 5% glycerol, 5 mM EDTA) supplemented with a complete Protease Inhibitor Cocktail tablet (Roche, Basel, Switzerland). Next, cells were lysed by sonication (50% duty cycle for 5 minutes with a 1-minute on/off cycle) and subsequently centrifuged at 30,000 rpm using a Ti70 rotor (Beckman Coulter Life Sciences, Indianapolis, IN) for 45 minutes at 4°C. Afterwards, the supernatant was loaded onto an Acta HPLC system (General Electric, Schenectady, NY) with a 1 mL nickel column for affinity chromatography. The bound protein was then eluted with elution buffer (20 mM HEPES (pH 7.5), 200 mM NaCl and 300 mM imidazole). The concentration of imidazole was reduced 40,000x by consecutive dialysis using an Invitrogen 3 ml dialysis cassette (ThermoFischer Scientific) in 20 mM HEPES (pH 7.5) and 200 mM NaCl solution. The remaining volume was concentrated to 1.5 ml using an Amicon Ultra cellulose 10,000 MWCO filtration unit (Millipore Sigma). The final protein concentration was quantified with a Pierce BCA assay (ThermoFischer Scientific) to be 2 mg/ml. Finally, 10% glycerol was added for storage at -80°C.

### Pronase digestion assay

Lyophilized *S. griseus* pronase (Roche) was dissolved in milli-Q water to make a stock concentration of 6 mg/mL, which was then stored at 4°C for up to three days. Following ANM generation, beads were pelleted by centrifugation at 300 xg for 5 minutes at 4°C. The supernatant was then carefully removed without disturbing the bead pellet, which was subsequently washed twice in 1 mL PBS (Ca^2+^ and Mg^2+^-free, pH 7.5) followed by resuspension in 30 μl PBS. Next, 18 μl of sample was loaded via capillary action into an imaging flow chamber made from a glass coverslip adhered to a glass slide, both of which were purchased from ThermoFischer Scientific, and separated by two strips of double-sided tape. Finally, 9 μl of pronase stock solution was added to the chamber containing the ANMs resulting in a final pronase concentration of 2 mg/mL. For the digestion of His_6_-GST-EGFP, purified protein was added to SUPER templates containing 60% DOPC, 30% cholesterol, and 10% Ni-NTA lipids at a final concentration of 1 μM, incubated at room temperature for 15 minutes, washed twice with PBS, and then resuspended in 30 μl PBS. Confocal images were acquired before and 15 minutes after the addition of pronase.

### KASH-binding assay

TRITC-KASH2^WT^ (SEDDYSCQANNFARSFYPMLRYTNGPPPT) and TRITC-KASH2^ΔPPPT^ (SEDDYSCQANNFARSFYPMLRYTNG) were purchased from Genscript Biotech (Piscataway, NJ). The lyophilized peptides were dissolved in DMSO to give a final stock concentration of 10 μM each. Similar to the pronase digestion assay, the ANMs were washed and resuspended in 30 μL PBS and then 10 μl aliquots were made for TRITC-KASH2 peptide binding tests. To each aliquot of washed ANMs, we added 0.5 μL of TRITC-KASH^WT^ or KASH2^ΔPPPT^ peptides as well as 0.5 μL milli-Q water resulting in a final peptide concentration of 450 nM. To each aliquot of washed SUPER templates, which were resuspended completely before being equally distributed in different tubes, we added 1.0 μL of TRITC-KASH^WT^ peptide resulting in a final peptide concentration of 1 μM. The binding reactions were then incubated for 15 minutes at room temperature prior to imaging.

### Microscopy

All images were acquired using an oil immersion 100X/1.4 NA Plan-Apochromat objective with an Olympus IX-81 inverted fluorescence microscope (Olympus Corporation, Tokyo, Japan) controlled by MetaMorph software (Molecular Devices, San Jose, CA) equipped with a CSU-X1 spinning disk confocal head (Yokogawa Electric Corporation, Tokyo, Japan), AOTF-controlled solid-state lasers (Andor Technology, Belfast, UK), and an iXON3 EMCCD camera (Andor Technology, Belfast, UK). Images of EGFP fluorescence images were acquired with 488 nm laser excitation at an exposure of 500 ms for all experimental conditions. TRITC fluorescence images were acquired with 561 nm laser excitation at an exposure of 100 ms. A Semrock 25 nm quad-band band-pass filter (FF01-440/521/607/700-25, IDEX Health and Science LLC, Rochester, NY) centered at 440, 521, 607, and 700 nm was used as the emission filter. Each acquired image contained ~3-5 beads, SUPER templates, or ANMs that had settled down upon a coverslip. For an individual experiment, 4 images were taken at different locations across a coverslip. Each experiment was repeated three independent times using the same imaging procedure. Samples were always freshly prepared before each experiment and were never reused.

### Image analysis

All images were analyzed using FIJI. We did not exclude any data from our analyses nor did we utilize blinding. Since the fluorescent rings corresponding to all membrane associating proteins were not homogenous in intensity, 8-10 line-scans were performed through the center of each bead at multiple angles and the maximum intensities of those scans were recorded. These values were averaged over each bead to generate one data point in the box plots (marking the first and third quartile with the box and the median) shown. Averaged background intensity measurements were performed for each image, which were subsequently subtracted from the individual fluorescence intensities of all beads present in that image. Normalization was carried out with respect to the maximum background subtracted intensity of beads in a given channel corresponding to the cell-free expression of a given protein in the absence of pronase. For the plots quantifying KASH peptide binding, normalization was carried out with respect to the maximum intensity of the beads incubated with TRITC-KASH2^WT^ peptides. Since the fluorescence intensities measured for each sample displayed very little variation, the ~16-20 beads, SUPER templates, or ANMs analyzed per condition were sufficient to enable the detection of statistically significant effect of particular experimental manipulation. Statistical analysis was performed using a 2-tailed *t*-test with a significance level of 0.05, as the data shown in this work meets its assumptions for statistical significance. The variance is conserved between the individual groups of data that were compared using this statistical test.

## Figure Legends

**Figure S1: CFE synthesized EGFP-SUN1^FL^-His_6_ and EGFP-SUN1^FL^-His_6_ are inserted into ER-derived microsomes.** Representative images of CFE reactions expressing the indicated SUN protein construct and stained with DiI taken **A)** within the CFE reaction solution or **B)** at the coverslip. Scale bar: 5 μm.

**Figure S2: Rapid reduction of EGFP fluorescence on SUPER templates by pronase. A)** Representative images of purified His_6_-GST-EGFP incubated with SUPER templates containing 10% DOGS-NTA-Ni before (t= 0 min) or after (t = 5 min) the addition of pronase. Scale bar: 5 μm. **B)** Box plots depicting the relative fluorescence units (RFU) of EGFP quantified on ANMs before (t = 0 min) and after (t = 5 min) the addition of pronase for His_6_-GST-EGFP from 3 independent experiments (n = 16 beads per condition). *** *p* < 0.001.

**Figure S3: Topology of Penta-His-AF647-labeled EGFP-SUN1^FL^-His_6_ and EGFP-SUN2^FL^-His_6_ inserted into ANMs determined by pronase protection assay.** Representative images of Penta-His-AF647-labeled EGFP-SUN1^FL^-His_6_ and EGFP-SUN2^FL^-His_6_ inserted into ANMs taken before and 30 minutes after the addition of pronase. Scale bar: 5 μm.

